## Supplementary Figures and Tables for "Population-level genome-wide STR typing in *Plasmodium* species reveals higher resolution population structure and genetic diversity relative to SNP typing"

**S1 Fig.** Comparison with *P. falciparum* gel electrophoresis data. Bubble plots representing the GE allele calls are plotted against HipSTR's calls, where the left plot has been shifted such that the bottom leftmost point lies on the origin. In plot (b) points are coloured according to the region where samples originated from. The line represents y = x. It is important to note that in most of these plots, the Milne Bay samples are typically off the line y = x, indicating potential underlying issues with the GE calls for Milne Bay samples.

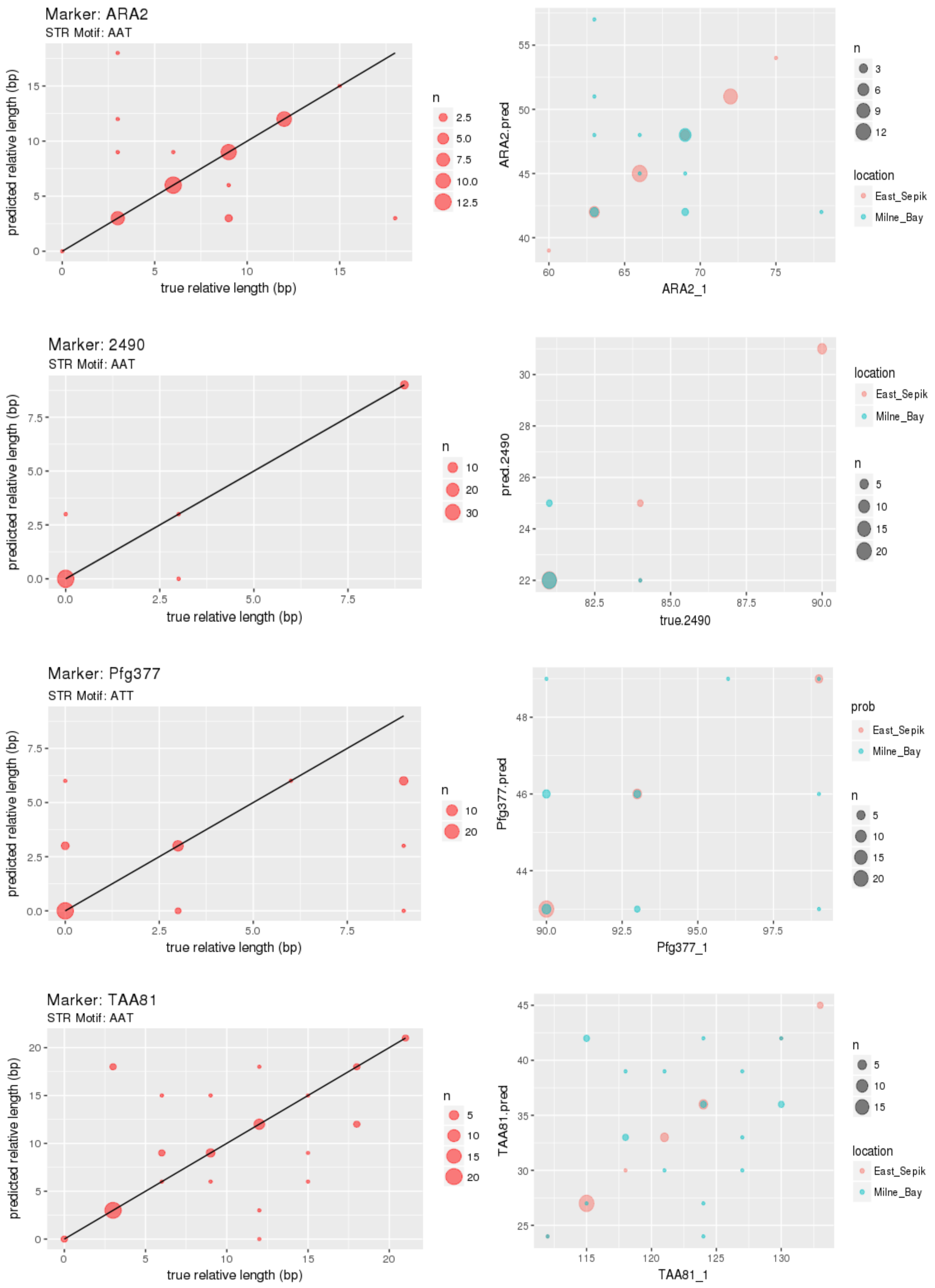

**S2 Fig.** ROC curves and AUC values of the *P. falciparum* complete dataset and five train datasets. (A) The mononucleotide STR model. (B) The polynucleotide STR model.

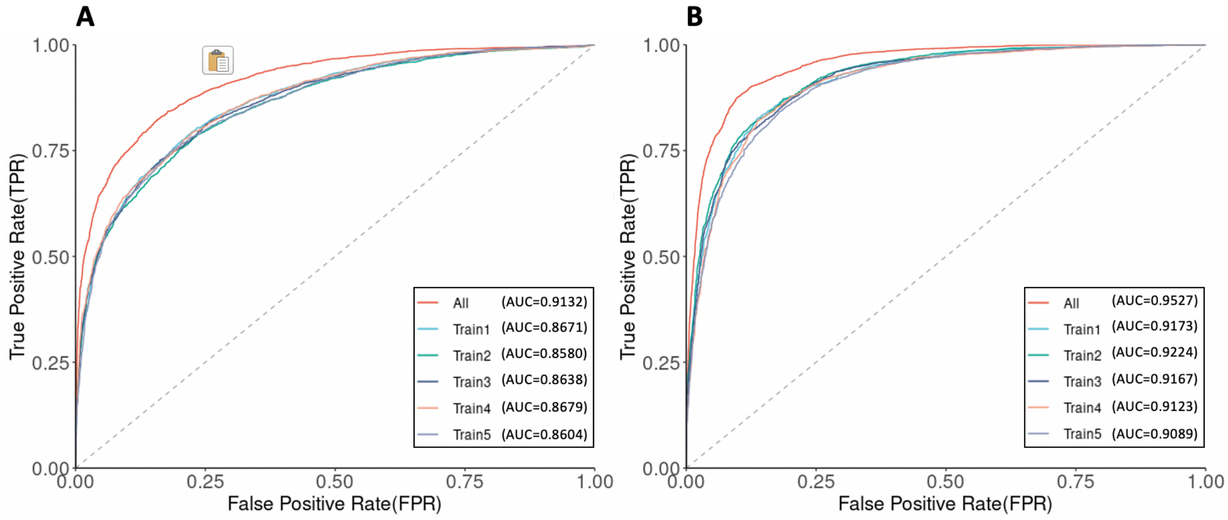

**S3 Fig.** AUC values of the *P. falciparum* fivefold cross validation-datasets (80/20, 70/20, 60/20, 50/20, 40/20, 30/20, 20/20, and 10/20 splits). (A) The mononucleotide STR model. (B) The polynucleotide STR model.

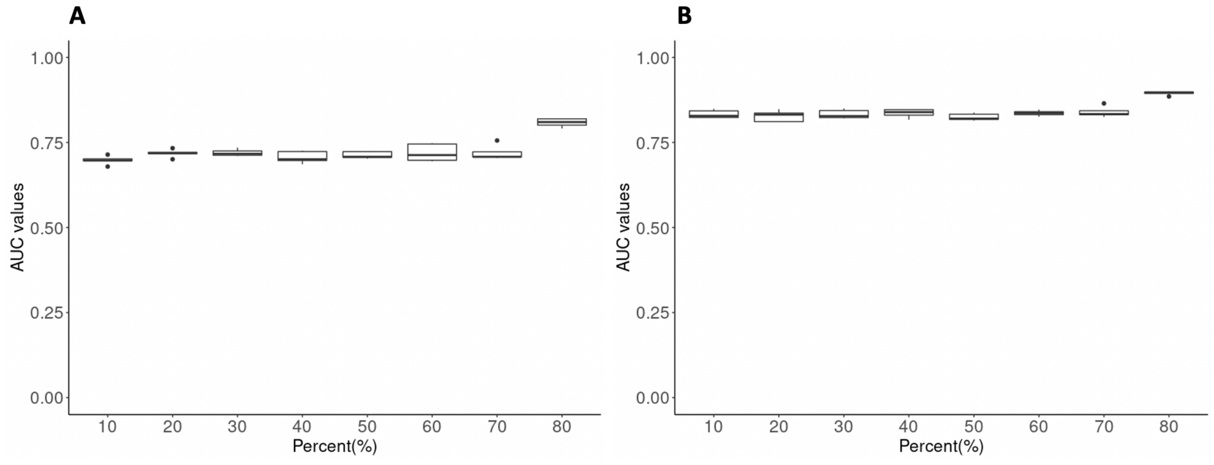

**S4 Fig.** Principal component analysis of the 3,047 *P. falciparum* samples of SNP and STR data. (A) SNP-based PCA based on 213,757 loci. (B) STR-based PCA based on 6,768 (2,563 mononucleotide STR and 4,205 polynucleotide STR) high-quality loci.

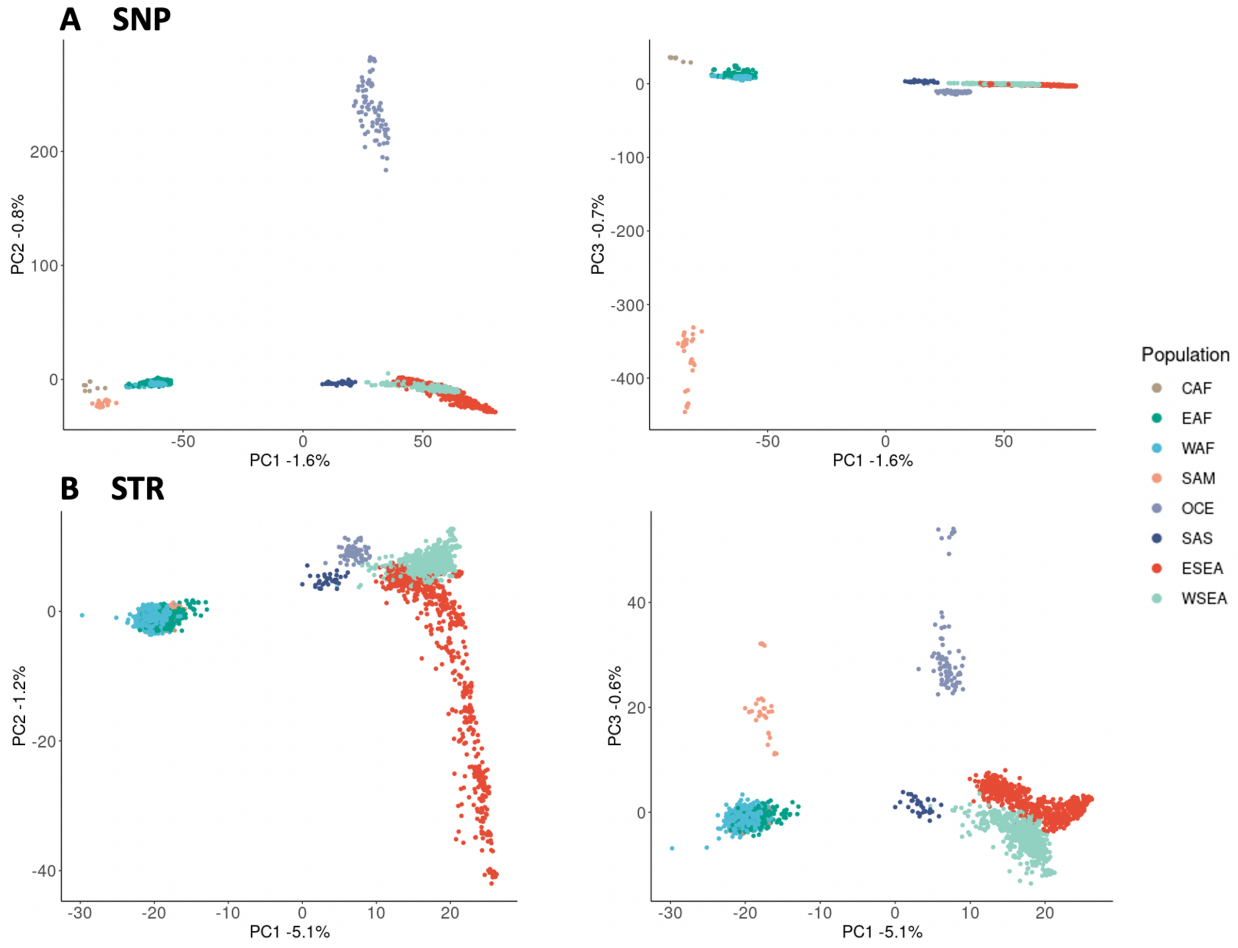

**S5 Fig.** Principal component analysis of the 174 *P. vivax* samples of SNP and STR data. (A) SNP-based PCA based on 188,571 loci. (B) STR-based PCA based on 3,496 (1,648 mononucleotide STR and 1,848 polynucleotide STR) high-quality loci.

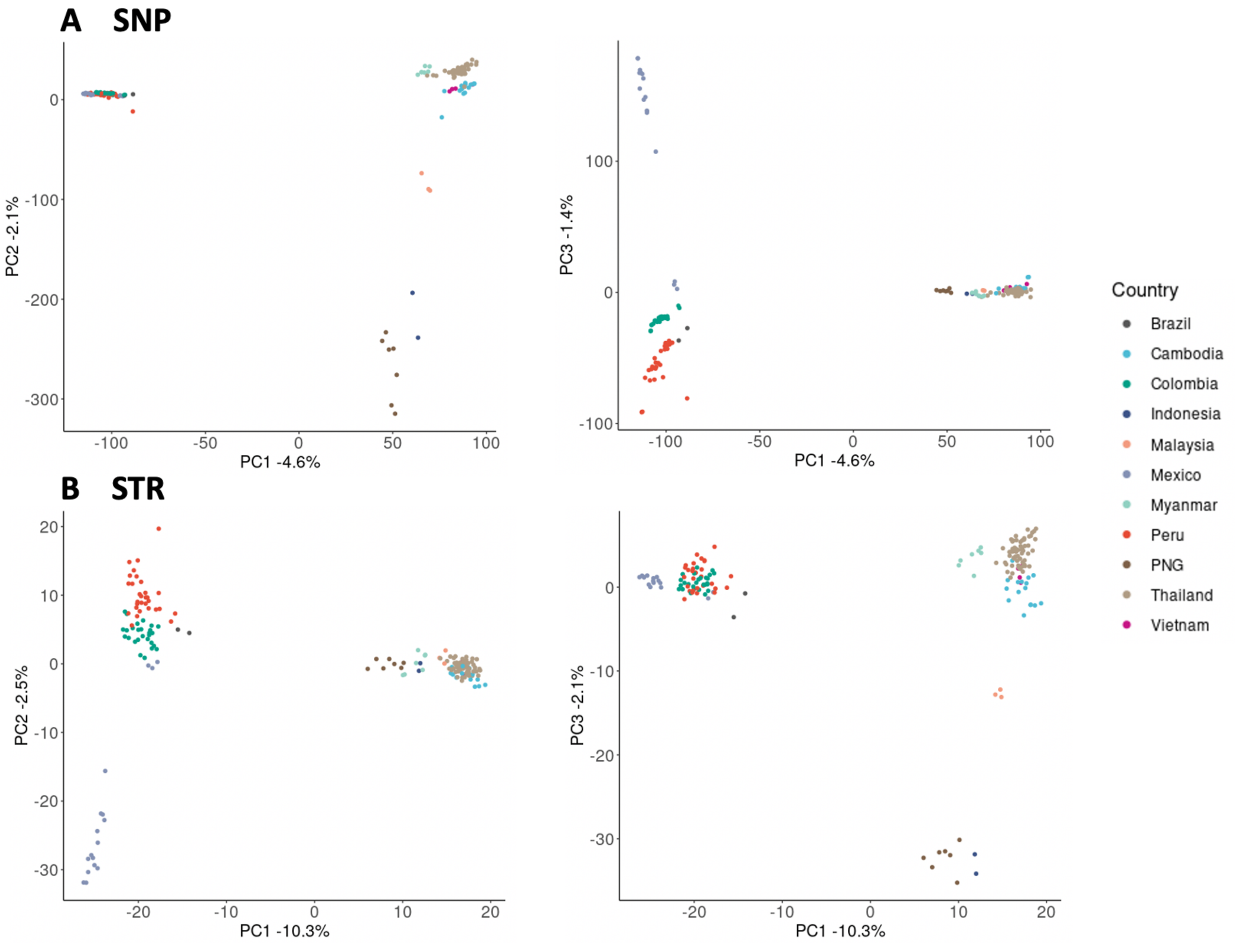

**S6 Fig.** Population structure analysis of the 174 *P. vivax* samples of SNP and STR data. (A) UMAP clustering of the top five principal components of the SNP data. (B) UMAP clustering of the top five principal components of the STR data with different colours representing the eight different countries. (C) NJTs based on the SNP data. (D) NJTs based on the STR data. Branches are colored according to the country.

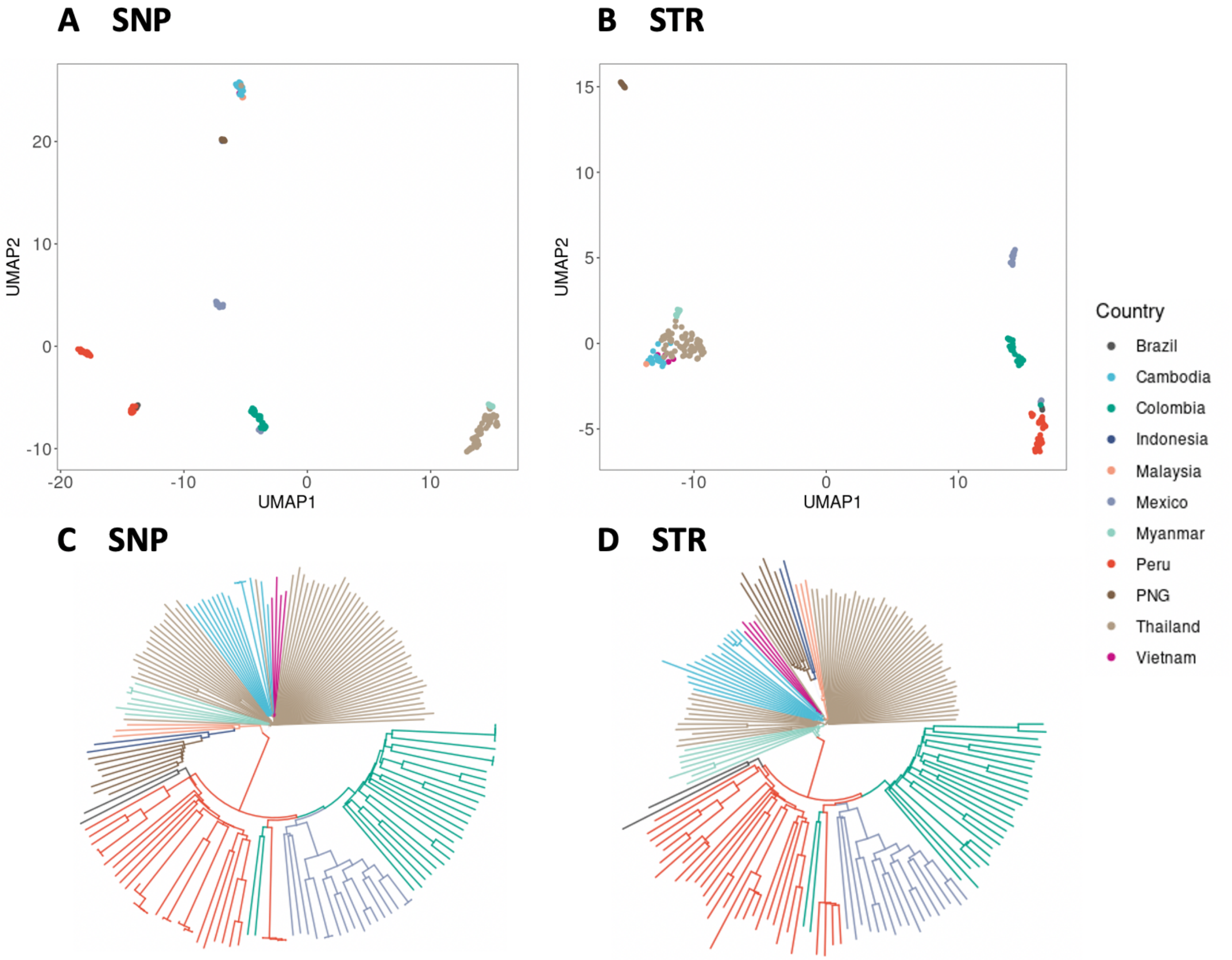

**S7 Fig.** Sub-population structure analysis of the SAM and EAF population *P. falciparum* samples of SNP and STR data. (A) UMAP on the top five principal components of the SNP data (SAM countries). (B) UMAP on the top five principal components of the STR data (SAM countries). Colouring the points by the SAM countries. (C) UMAP on the top five principal components of the SNP data (EAF countries). (D) UMAP on the top five principal components of the STR data (EAF countries). Colouring the points by the EAF countries.

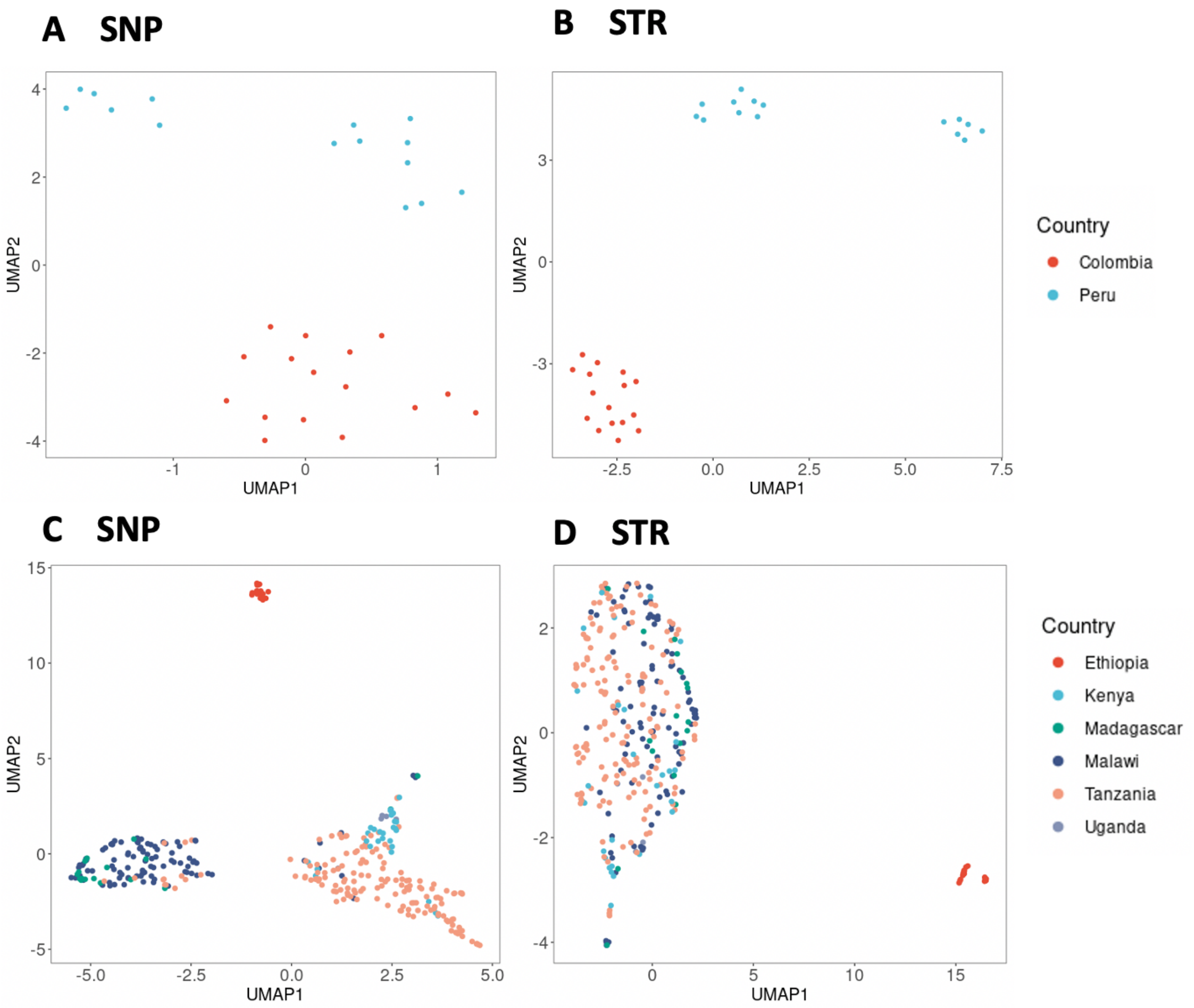

**S8 Fig.** A comparison of measures of genetic differentiation (*Jost's D* and $F_{ST}$) estimates using SNP and STR data of *P. falciparum*. (A) *Jost's D* of population pairs (*Mantel r* = 0.996, *P* = 0.001). (B) *Jost's D* of country pairs (*Mantel r* = 0.996, *P* = 0.001). (C) $F_{ST}$ of population pairs (*Mantel r* = 0.97, *P* = 0.001). (D) $F_{ST}$ of country pairs (*Mantel r* = 0.90, *P* = 0.001). Mantel tests were used to measure the correlation.

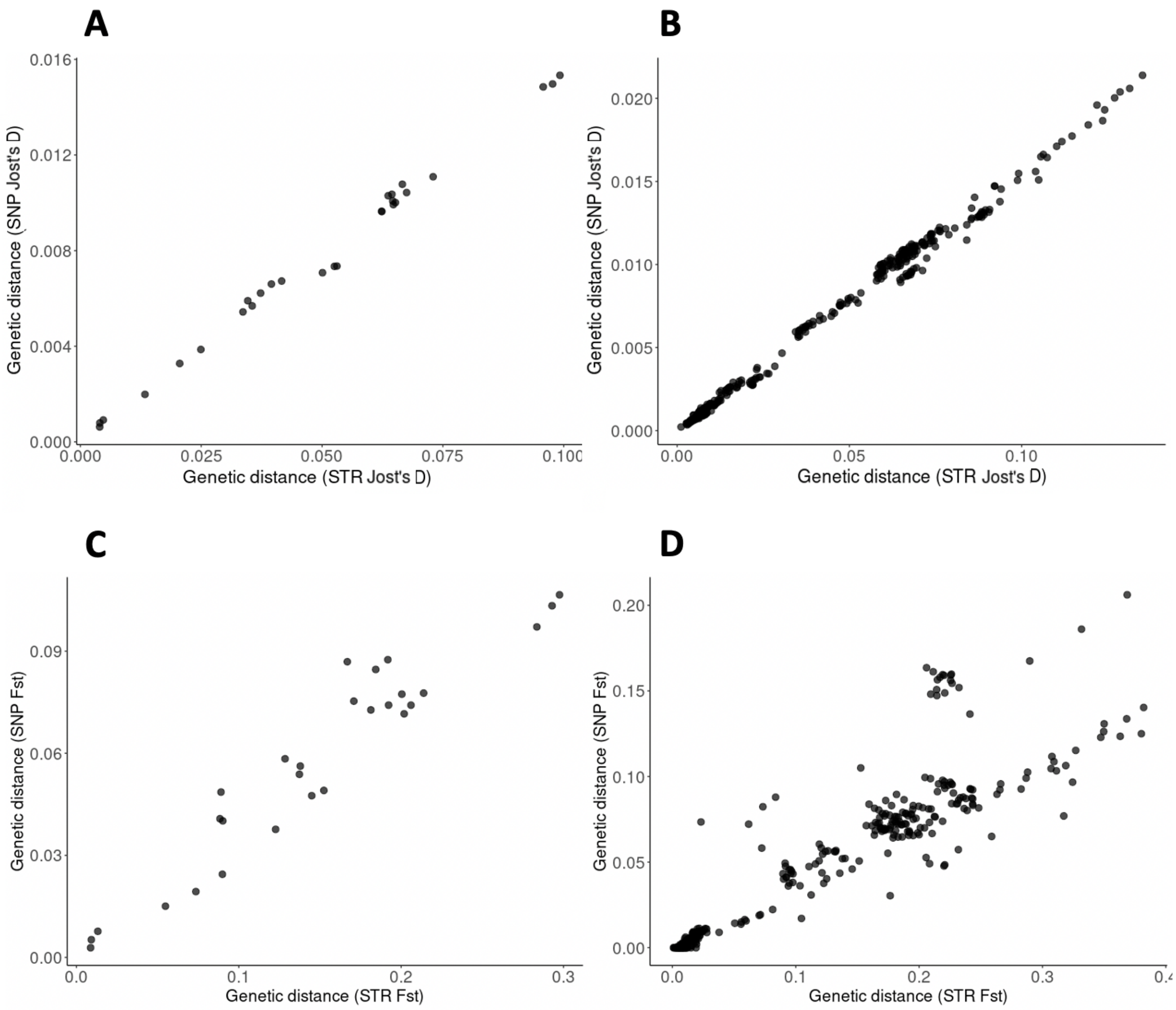

**S9 Fig.** A comparison of measures of genetic differentiation (*Jost's D* and $F_{ST}$) estimates using SNP and STR data of *P. vivax*. (A) *Jost's D* of country pairs (*Mantel r* = 0.9678, *P* = 0.001). (B) $F_{ST}$ of country pairs (*Mantel r* = 0.9185, *P* = 0.001). Mantel tests were used to measure the correlation.

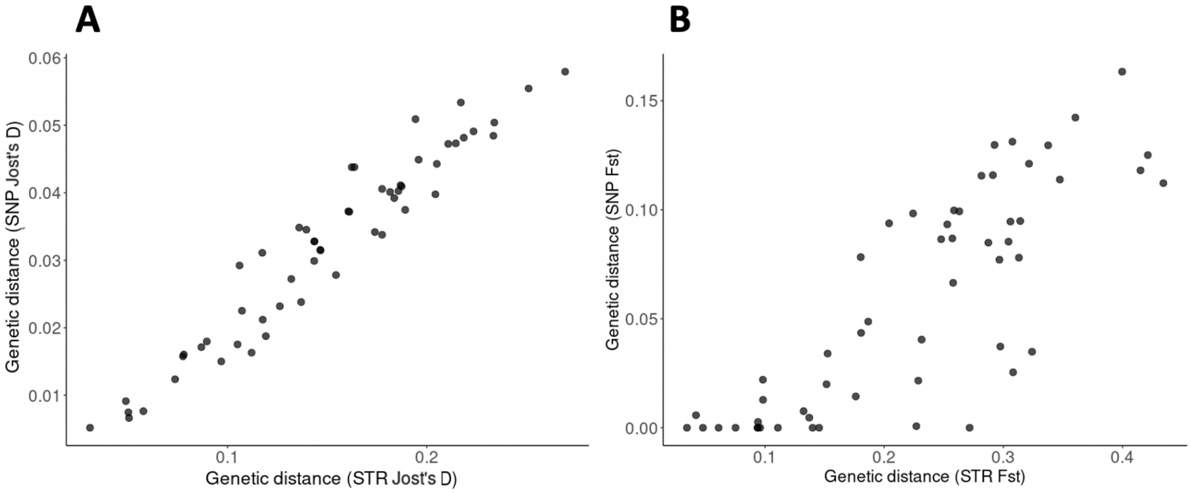

**S10 Fig.** Pairwise genetic distance (*Jost's D*) and geographical distances (km) between populations and countries of *P. falciparum*. (A) SNP data of populations. (B) STR data of populations. (C) SNP data of countries. (D) SNP data of countries. A Mantel test was used to measure the association.

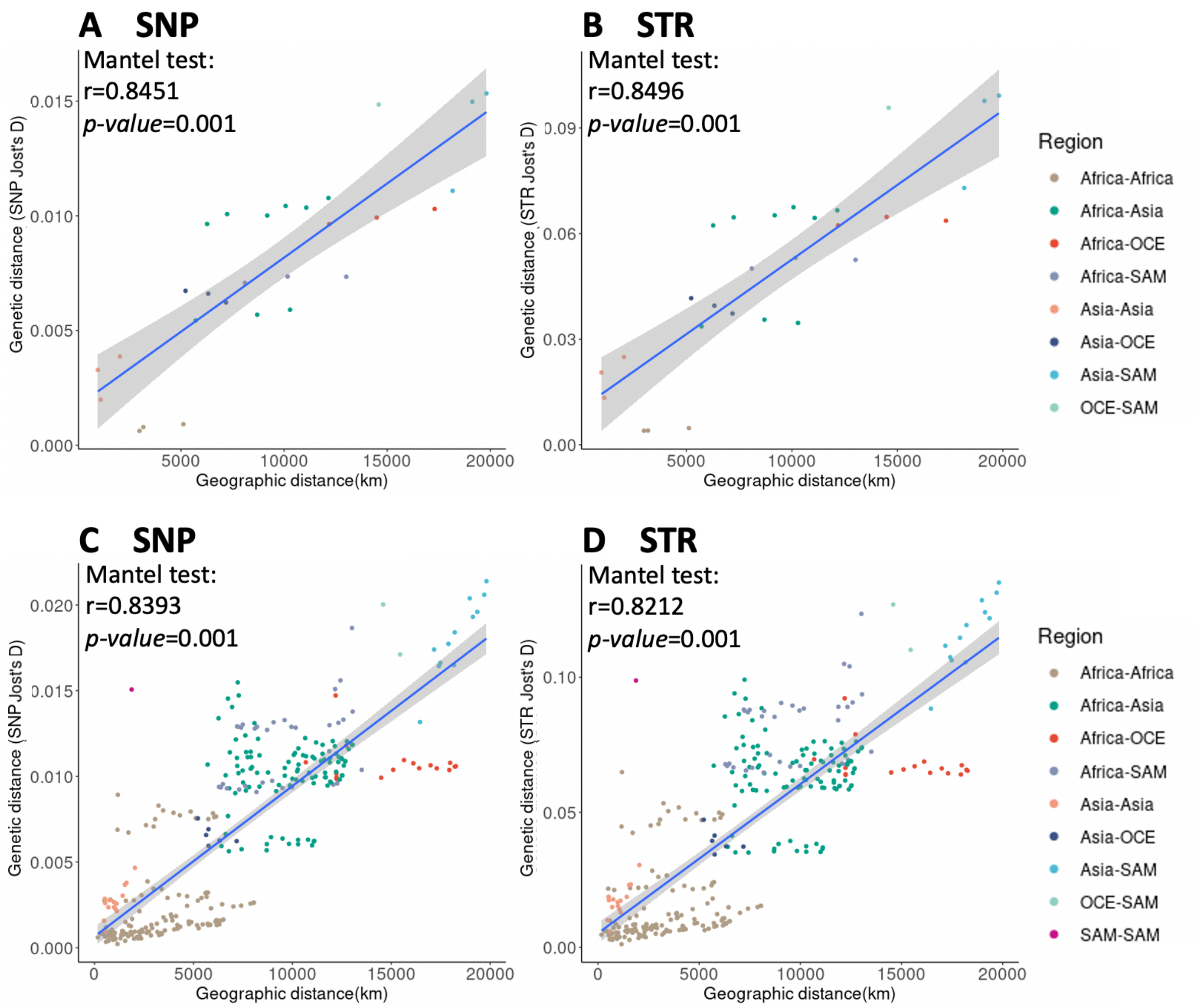

**S11 Fig.** Pairwise genetic distance (*Jost's D* and $F_{ST}$) and geographical distances (km) between countries of *P. vivax*. (A) SNP data of country pairs (*Jost's D*). (B) STR data of country pairs (*Jost's D*). (C) SNP data of country pairs ($F_{ST}$). (D) SNP data of country pairs ($F_{ST}$). A Mantel test was used to measure the association.

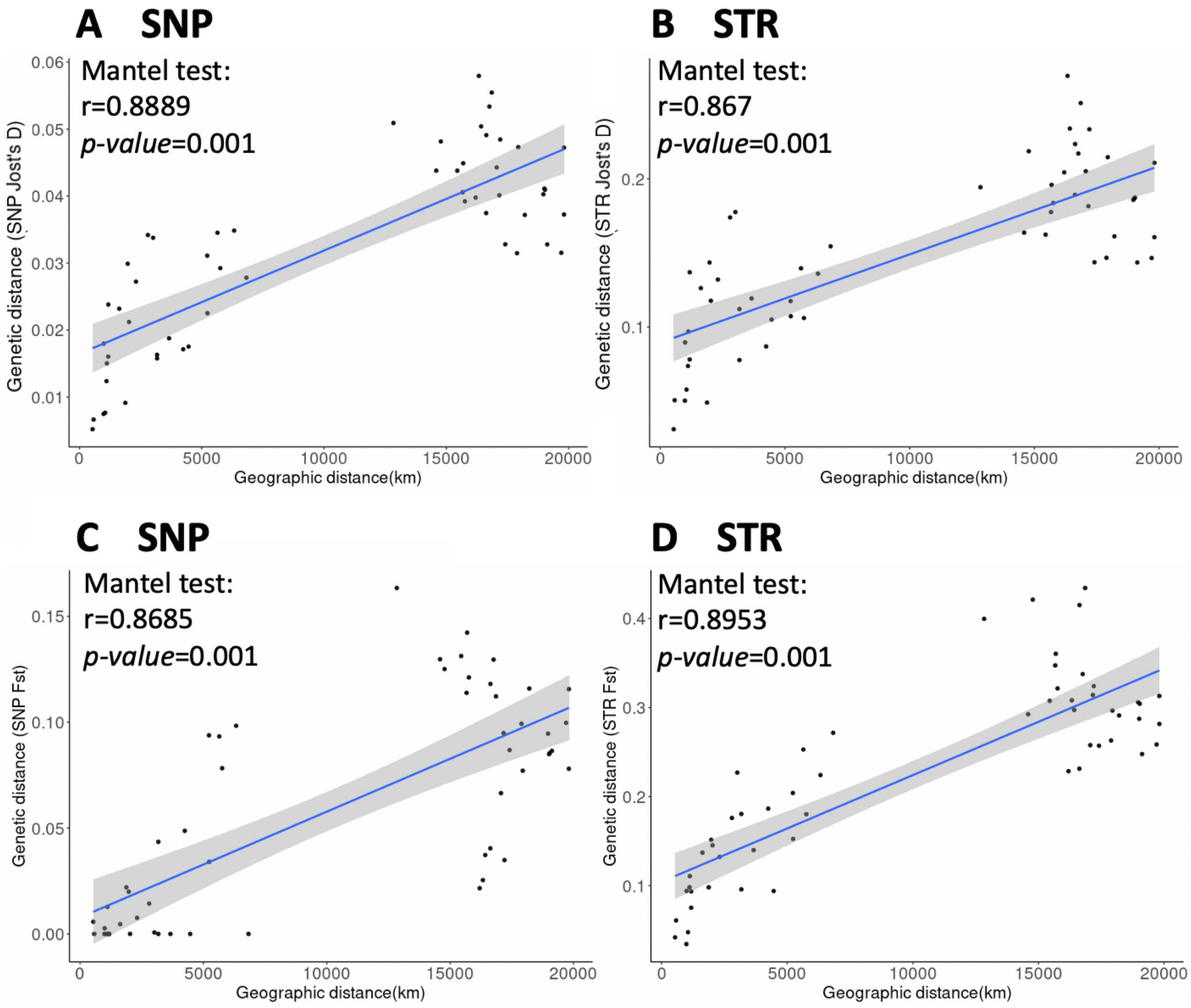

**S12 Fig.** Minimum spanning network using Bruvo's distances based on the ten most informative STR markers showing the relationship among two groups of *P.* vivax isolates. (A) Cambodia and Thailand. (B) Colombia and Peru. (C) Mexico and Peru. Colors correspond to the country. Node sizes correspond to the number of samples. Edge lengths are arbitrary.

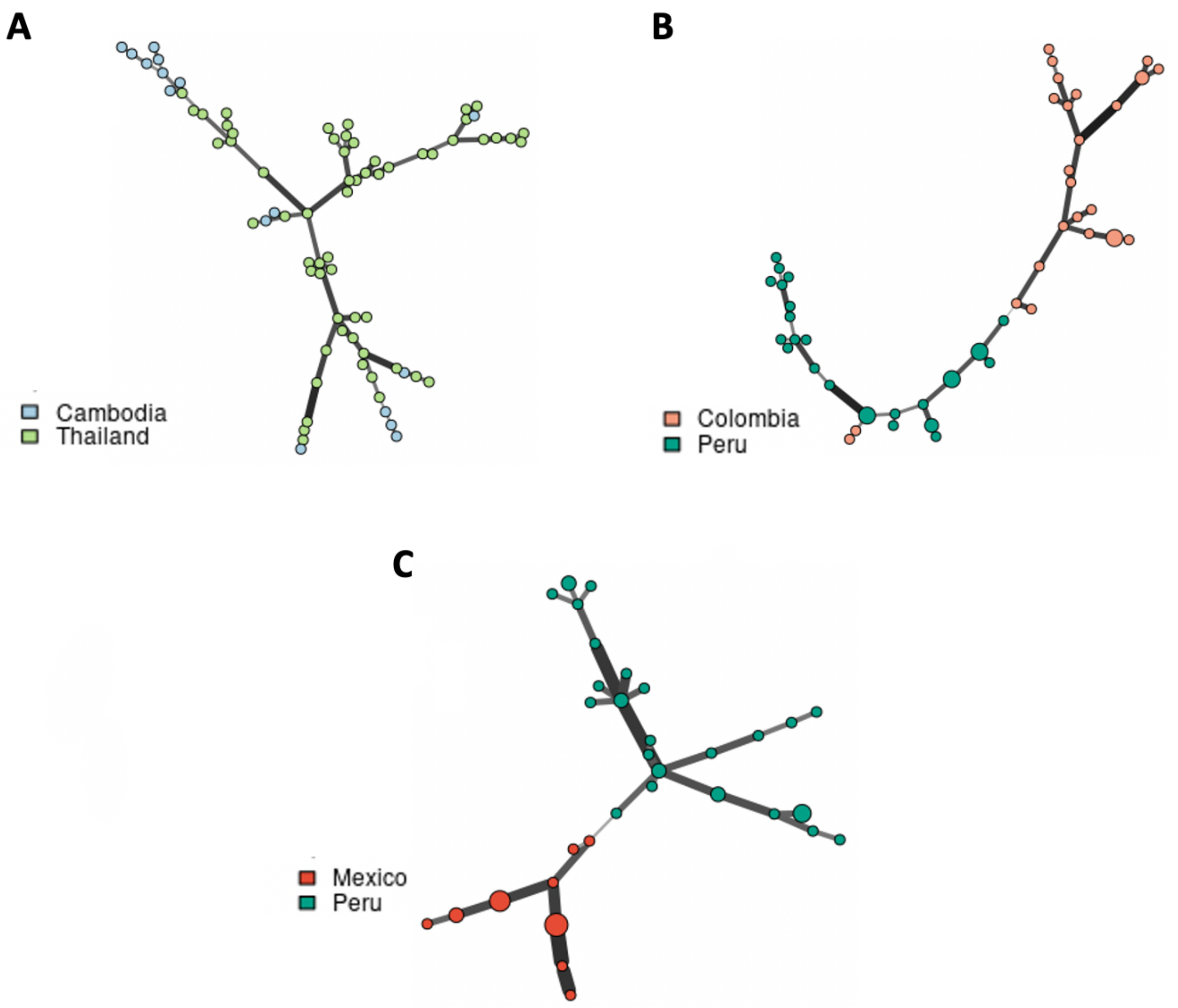

**S13 Fig.** Examples of corresponding genotypes. (A) One drug-resistant sample. (B) One drug-sensitive sample.

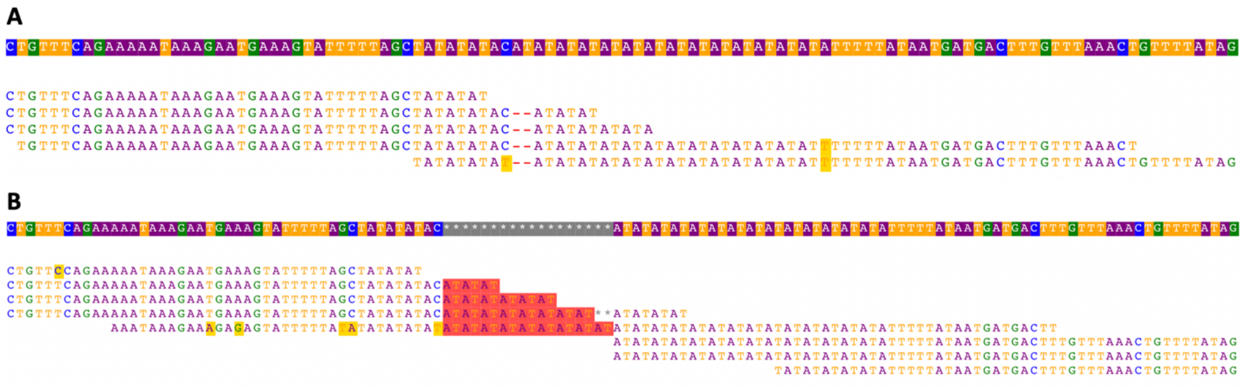

### Supplementary tables

**S1 Table.** Baseline variables used for prediction of the STR quality.

| Variable | Description |
|---|---|
| **STR feature** | |
| Length | Motif size (1-9bp) |
| Repeat units | Number of times the motif is repeated in tandem |
| GC_STR | GC content of STR regions |
| GC_Flank | GC content of 100-bp-long STR flanking regions (50 bp each side) |
| GC_Diff | GC content difference between the STR regions and STR flanking regions |
| Sequence_Complexity | Sequence complexity of the repeat regions |
| Missingness | Missingness of the STR genotype |
| **HipSTR parameters (Posterior, Stutter, Indel)** | |
| Mean_Posterior | The mean posterior probability of the STR genotype across all samples |
| Mean_Indel | The mean total number of reads across all samples contains an indel in the regions flanking the STR |
| Mean_Stutter | The mean number of reads across all samples at a locus with a stutter artifact |
| **STR features that were associated with population** | |
| ***P. falciparum*** **dataset is based on Population-level labels,** ***P. vivax*** **dataset is based on Country-level labels** | |
| He | The expected heterozygosity, a measure of genetic variation within populations |
| MeanHe | Mean heterozygosity of different populations |
| MinimumHe | Minimum heterozygosity of different populations |
| MaximumHe | Maximum heterozygosity of different populations |
| JostD | The degree of population differentiation measured by *Jost's D* |

**S2 Table.** Multivariable logistic regression model's estimated coefficients and respective 95% confidence intervals. The model was fitted on the *P. falciparum* dataset without the population origin label.

| | Variable | Coefficient estimate | Lower 95% CI | Upper 95% CI | *P* values |
|---|---|---|---|---|---|
| **Modeling Mononucleotide STR** | (Intercept) | 0.24 | 0.11 | 0.42 | 0.002 |
| | Repeat units | 0.11 | 0.04 | 0.19 | 0.002 |
| | GC_Diff | 0.10 | 0.03 | 0.17 | 0.004 |
| | Mean_Indel | 0.08 | -0.02 | 0.18 | 0.1 |
| | Mean_Posterior | 0.65 | 0.54 | 0.75 | <0.001 |
| | Mean_stutter | -0.21 | -0.33 | -0.10 | <0.001 |
| | He | 7.55 | 6.67 | 8.43 | <0.001 |
| | MeanHe | -5.57 | -6.57 | -4.56 | <0.001 |
| | JostD | -1.16 | -1.66 | -0.40 | <0.001 |
| | MinimumHe | -0.60 | -0.81 | -0.39 | <0.001 |
| | MaximumHe | 3.39 | 3.08 | 3.71 | <0.001 |
| **Modeling Polynucleotide STR** | (Intercept) | 1.74 | 1.58 | 1.90 | <0.001 |
| | Length | -0.13 | -0.20 | -0.05 | <0.001 |
| | Repeat units | 0.16 | 0.06 | 0.25 | 0.001 |
| | GC_Flank | 0.09 | 0.03 | 0.15 | 0.003 |
| | Mean_Posterior | 0.09 | -0.02 | 0.18 | 0.09 |
| | Mean_Stutter | -0.14 | -0.22 | -0.01 | 0.004 |
| | He | 10.55 | 8.87 | 12.23 | <0.001 |
| | MeanHe | -16.27 | -18.04 | -14.51 | <0.001 |
| | JostD | -0.76 | -1.14 | -0.37 | <0.001 |
| | MinimumHe | 1.16 | 0.85 | 1.47 | <0.001 |
| | MaximumHe | 9.18 | 8.64 | 9.73 | <0.001 |

**S3 Table.** Multivariable logistic regression model's estimated coefficients and respective 95% confidence intervals. The model was fitted on the *P. vivax* dataset.

| | **Variable** | **Coefficient estimate** | **Lower 95% CI** | **Upper 95% CI** | ***P* values** |
|---|---|---|---|---|---|
| **Modeling Mononucleotide STR** | (Intercept) | -2.13 | -2.21 | -2.06 | <0.001 |
| | Repeat units | -0.06 | -0.14 | 0.01 | 0.09 |
| | GC_STR | -0.15 | -0.21 | -0.10 | <0.001 |
| | GC_Flank | 0.14 | -0.02 | 0.30 | 0.09 |
| | GC_Diff | -0.12 | -0.28 | 0.04 | 0.15 |
| | Mean_Posterior | 0.66 | 0.59 | 0.74 | <0.001 |
| | Mean_stutter | -0.17 | -0.25 | -0.09 | <0.001 |
| | He | 3.79 | 3.61 | 3.98 | <0.001 |
| | MeanHe | -2.69 | -2.90 | -2.48 | <0.001 |
| | MinimumHe | 0.08 | 0.04 | 0.13 | <0.001 |
| | MaximumHe | 1.42 | 1.31 | 1.53 | <0.001 |
| **Modeling Polynucleotide STR** | (Intercept) | -2.88 | -3.01 | -2.75 | <0.001 |
| | Repeat units | -0.06 | -0.13 | 0.02 | 0.12 |
| | Missingness | 0.08 | 0.00 | 0.15 | 0.06 |
| | Mean_Posterior | -0.08 | -0.15 | -0.01 | 0.03 |
| | He | 3.37 | 3.10 | 3.64 | <0.001 |
| | MeanHe | -3.10 | -3.40 | -2.81 | <0.001 |
| | JostD | -0.10 | -0.21 | 0.00 | 0.06 |
| | MinimumHe | 0.14 | 0.08 | 0.20 | <0.001 |
| | MaximumHe | 1.86 | 1.72 | 2.00 | <0.001 |

**S4 Table.** Multivariable logistic regression model's estimated coefficients and respective 95% confidence intervals. The model was fitted on the *P. vivax* dataset without the population origin label.

| | **Variable** | **Coefficient estimate** | **Lower 95% CI** | **Upper 95% CI** | ***P* values** |
|---|---|---|---|---|---|
| **Modeling Mononucleotide STR** | (Intercept) | -2.20 | -2.28 | -2.13 | <0.001 |
| | Repeat units | -0.12 | -0.20 | -0.05 | 0.002 |
| | GC_STR | -0.24 | -0.29 | -0.18 | <0.001 |
| | GC_Flank | 0.26 | 0.09 | 0.43 | 0.002 |
| | GC_Diff | -0.23 | -0.40 | -0.06 | 0.007 |
| | Mean_Posterior | 0.59 | 0.52 | 0.67 | <0.001 |
| | Mean_stutter | -0.14 | -0.22 | -0.07 | <0.001 |
| | He | 2.34 | 2.12 | 2.56 | <0.001 |
| | MeanHe | -1.28 | -1.59 | -0.98 | <0.001 |
| | MinimumHe | -0.74 | -0.88 | -0.60 | <0.001 |
| | MaximumHe | 1.77 | 1.57 | 1.98 | <0.001 |
| | JostD | 0.33 | 0.25 | 0.42 | <0.001 |
| **Modeling Polynucleotide STR** | (Intercept) | -2.85 | -2.97 | -2.73 | <0.001 |
| | Repeat units | -0.20 | -0.28 | -0.12 | <0.001 |
| | Length | -0.13 | -0.22 | -0.05 | 0.002 |
| | GC_STR | -0.07 | -0.17 | 0.02 | 0.14 |
| | He | 0.71 | 0.46 | 0.95 | <0.001 |
| | MeanHe | -1.56 | -1.92 | -1.20 | <0.001 |
| | JostD | 0.07 | -0.02 | 0.16 | 0.15 |
| | MinimumHe | -0.16 | -0.32 | -0.01 | 0.04 |
| | MaximumHe | 2.94 | 2.66 | 3.22 | <0.001 |

**S5 Table.** Detected selection signatures (located in the coding region) between the *P. falciparum* populations containing the top 0.1% of STR.

| Gene | Product Description | Genomic Location | Population | JostD |
|---|---|---|---|---|
| PF3D7_0627800 | acetyl-CoA synthetase, putative | Pf3D7_06_v3:g.1116360ATT[12] | ESEA-SAM | 1 |
| PF3D7_0627800 | acetyl-CoA synthetase, putative | Pf3D7_06_v3:g.1116360ATT[12] | OCE-SAM | 1 |
| PF3D7_0627800 | acetyl-CoA synthetase, putative | Pf3D7_06_v3:g.1116360ATT[12] | SAS-SAM | 1 |
| PF3D7_0627800 | acetyl-CoA synthetase, putative | Pf3D7_06_v3:g.1116360ATT[12] | WSEA-SAM | 1 |
| PF3D7_0628100 | HECT-domain (ubiquitin-transferase), putative | Pf3D7_06_v3:g.1151898ATT[10] | ESEA-SAS | 0.79 |
| PF3D7_0628100 | HECT-domain (ubiquitin-transferase), putative | Pf3D7_06_v3:g.1125666ATT[6] | ESEA-SAS | 0.73 |
| PF3D7_0629700 | SET domain protein, putative | Pf3D7_06_v3:g.1231112AAT[9] | CAF-SAM | 0.99 |
| PF3D7_0629700 | SET domain protein, putative | Pf3D7_06_v3:g.1231112AAT[9] | EAF-SAM | 0.97 |
| PF3D7_0629700 | SET domain protein, putative | Pf3D7_06_v3:g.1231112AAT[9] | ESEA-OCE | 0.93 |
| PF3D7_0629700 | SET domain protein, putative | Pf3D7_06_v3:g.1231112AAT[9] | ESEA-SAM | 1 |
| PF3D7_0629700 | SET domain protein, putative | Pf3D7_06_v3:g.1231112AAT[9] | ESEA-SAS | 0.88 |
| PF3D7_0629700 | SET domain protein, putative | Pf3D7_06_v3:g.1232864AATAATGTG[4] | ESEA-SAS | 0.72 |
| PF3D7_0629700 | SET domain protein, putative | Pf3D7_06_v3:g.1231112AAT[9] | ESEA-WSEA | 0.54 |
| PF3D7_0629700 | SET domain protein, putative | Pf3D7_06_v3:g.1232864AATAATGTG[4] | ESEA-WSEA | 0.54 |
| PF3D7_0629700 | SET domain protein, putative | Pf3D7_06_v3:g.1231112AAT[9] | OCE-SAM | 1 |
| PF3D7_0629700 | SET domain protein, putative | Pf3D7_06_v3:g.1231112AAT[9] | SAS-SAM | 1 |
| PF3D7_0629700 | SET domain protein, putative | Pf3D7_06_v3:g.1231112AAT[9] | WAF-SAM | 0.93 |
| PF3D7_0629700 | SET domain protein, putative | Pf3D7_06_v3:g.1227974AAT[8] | WSEA-SAM | 1 |
| PF3D7_0709300 | Cg2 protein | Pf3D7_07_v3:g.417234ATT[6] | WSEA-OCE | 0.9 |
| PF3D7_0723900 | RNA-binding protein, putative | Pf3D7_07_v3:g.1003328ATT[9] | WSEA-SAM | 1 |
| PF3D7_0810600 | ATP-dependent RNA helicase DBP1, putative | Pf3D7_08_v3:g.544455AAT[9] | CAF-EAF | 0.72 |
| PF3D7_0810600 | ATP-dependent RNA helicase DBP1, putative | Pf3D7_08_v3:g.544455AAT[9] | CAF-SAS | 0.84 |
| PF3D7_0810600 | ATP-dependent RNA helicase DBP1, putative | Pf3D7_08_v3:g.544455AAT[9] | EAF-WAF | 0.79 |
| PF3D7_0810600 | ATP-dependent RNA helicase DBP1, putative | Pf3D7_08_v3:g.544455AAT[9] | WAF-SAS | 0.87 |
| PF3D7_0810900 | conserved Plasmodium protein, unknown function | Pf3D7_08_v3:g.551208ATTTTC[4] | CAF-EAF | 0.74 |
| PF3D7_0810900 | conserved Plasmodium protein, unknown function | Pf3D7_08_v3:g.551208ATTTTC[4] | EAF-WAF | 0.7 |
| PF3D7_0811200 | ER membrane protein complex subunit 1, putative | Pf3D7_08_v3:g.562938AAT[11] | CAF-EAF | 0.6 |
| PF3D7_0811200 | ER membrane protein complex subunit 1, putative | Pf3D7_08_v3:g.562938AAT[11] | CAF-WAF | 0.47 |
| PF3D7_0826100 | HECT-like E3 ubiquitin ligase, putative | Pf3D7_08_v3:g.1121887ATT[19] | ESEA-WSEA | 0.47 |
| PF3D7_0916400 | conserved Plasmodium protein, unknown function | Pf3D7_09_v3:g.687263AATACT[4] | CAF-ESEA | 0.97 |
| PF3D7_0916400 | conserved Plasmodium protein, unknown function | Pf3D7_09_v3:g.687263AATACT[4] | EAF-ESEA | 0.97 |
| PF3D7_1107300 | polyadenylate-binding protein-interacting protein 1, putative | Pf3D7_11_v3:g.303973ATT[9] | CAF-SAS | 0.81 |
| PF3D7_1107300 | polyadenylate-binding protein-interacting protein 1, putative | Pf3D7_11_v3:g.303973ATT[9] | EAF-SAS | 0.74 |
| PF3D7_1208200 | cysteine repeat modular protein 3 | Pf3D7_12_v3:g.380602AAT[8] | CAF-OCE | 0.94 |
| PF3D7_1208200 | cysteine repeat modular protein 4 | Pf3D7_12_v3:g.380602AAT[8] | EAF-OCE | 0.97 |
| PF3D7_1208200 | cysteine repeat modular protein 5 | Pf3D7_12_v3:g.380602AAT[8] | WAF-OCE | 0.95 |
| PF3D7_1221000 | histone-lysine N-methyltransferase, H3 lysine-4 specific | Pf3D7_12_v3:g.843354ATT[9] | EAF-WSEA | 0.93 |
| PF3D7_1224700 | conserved Plasmodium protein, unknown function | Pf3D7_12_v3:g.1006995ATT[6] | SAS-WSEA | 0.59 |

| | | | | |
|---|---|---|---|---|
| PF3D7_1317900 | nucleolar complex protein 4, putative | Pf3D7_13_v3:g.744342ATT[8] | SAS-WSEA | 0.8 |
| PF3D7_1319900 | conserved Plasmodium protein, unknown function | Pf3D7_13_v3:g.824035ATC[4] | EAF-SAS | 0.75 |
| PF3D7_1336700 | parasitophorous vacuolar protein 3, putative | Pf3D7_13_v3:g.1482619ATT[7] | SAS-OCE | 0.87 |
| PF3D7_1431400 | surface-related antigen SRA | Pf3D7_14_v3:g.1234853T[13] | EAF-ESEA | 0.96 |
| PF3D7_1431400 | surface-related antigen SRA | Pf3D7_14_v3:g.1234853T[13] | WAF-ESEA | 0.97 |
| PF3D7_1431400 | surface-related antigen SRA | Pf3D7_14_v3:g.1234853T[13] | WAF-WSEA | 0.94 |
| PF3D7_1453600 | RAP protein, putative | Pf3D7_14_v3:g.2201945AT[8] | CAF-ESEA | 0.96 |
| PF3D7_1453600 | RAP protein, putative | Pf3D7_14_v3:g.2201945AT[8] | WAF-ESEA | 0.96 |
| PF3D7_1455300 | conserved protein, unknown function | Pf3D7_14_v3:g.2260430AAT[7] | CAF-WSEA | 0.95 |
| PF3D7_1455300 | conserved protein, unknown function | Pf3D7_14_v3:g.2260430AAT[7] | EAF-ESEA | 0.96 |
| PF3D7_1455300 | conserved protein, unknown function | Pf3D7_14_v3:g.2260430AAT[7] | EAF-WSEA | 0.97 |
| PF3D7_1455300 | conserved protein, unknown function | Pf3D7_14_v3:g.2260430AAT[7] | WAF-ESEA | 0.96 |
| PF3D7_1455300 | conserved protein, unknown function | Pf3D7_14_v3:g.2260430AAT[7] | WAF-WSEA | 0.97 |

**S6 Table.** Detected selection signatures (located in the coding region) between the *P. vivax* countries containing the top 0.1% of STR.

| Gene | Product Description | Genomic Location | Country | JostD |
|---|---|---|---|---|
| PVP01_0107900 | E3 ubiquitin-protein ligase, putative | PvP01_01_v1:g.396926AGCGGGGGT[4] | Cambodia-Colombia | 1 |
| PVP01_0107900 | E3 ubiquitin-protein ligase, putative | PvP01_01_v1:g.396926AGCGGGGGT[4] | Cambodia-Mexico | 1 |
| PVP01_0107900 | E3 ubiquitin-protein ligase, putative | PvP01_01_v1:g.396926AGCGGGGGT[4] | Cambodia-Peru | 1 |
| PVP01_0527500 | protein SOC1, putative | PvP01_05_v1:g.1138266CTT[7] | Cambodia-Colombia | 1 |
| PVP01_0527700 | conserved Plasmodium protein, unknown function | PvP01_05_v1:g.1145237CCT[9] | Peru-Colombia | 0.92 |
| PVP01_0709800 | cysteine repeat modular protein 1, putative | PvP01_07_v1:g.498494ACCGTTGAG[6] | Thailand-Colombia | 1 |
| PVP01_0709800 | cysteine repeat modular protein 1, putative | PvP01_07_v1:g.498494ACCGTTGAG[6] | Thailand-Mexico | 1 |
| PVP01_0834400 | P-loop containing nucleoside triphosphate hydrolase, putative | PvP01_08_v1:g.1449968CTT[5] | Colombia-Mexico | 0.99 |

**S7 Table.** *P. falciparum* STR markers used in the analysis and the number of samples genotyped at each STR marker. The symbol "*" denotes markers which were dropped in the TRF post-processing phase rather than the HipSTR phase. The table also highlights the location of the markers on the Pf3D7 reference genome as determined by the BLAST search.

| **STR Marker** | **Samples** | **Location** |
|---|---|---|
| TA1 | 0 | Chr06: 899846-900037 |
| Polyα | 0 | Chr04: 532133-532332 |
| TAA81 | 90 | Chr05: 1214321-1214442 |
| TA60 | 0* | Chr13: 2588612-2588932 |
| ARA2 | 84 | Chr11: 416197-416367 |
| Pfg377 | 85 | Chr12: 2045792-2045921 |
| PfPK2 | 0 | Chr12: 1611149-1611390 |
| TAA87 | 82 | Chr06: 374731-374830 |
| TAA42 | 0* | Chr05: 325646-325885 |
| 2490 | 87 | Chr10: 458210-458349 |
